# A microcircuit model of astrocytic potassium buffering and neural synchronization

**DOI:** 10.64898/2026.06.15.732376

**Authors:** Marco Cafiso, Gabriele Casagrande, Marianna Angiolleli, Paolo Paradisi, Pierpaolo Sorrentino, Damien Depannemaecker

## Abstract

Neural synchronization is fundamental to brain function and, when it becomes excessive, underlies pathological conditions such as epilepsy. Among brain regions, the temporal lobes, and the hippocampus in particular, exhibit the highest epileptogenic potential, with mesial temporal lobe epilepsy representing the most prevalent form of the condition in humans. Within the hippocampus, extracellular potassium dynamics are central to non-synaptic epileptiform activity, and astrocytic potassium buffering mechanisms have emerged as key regulators of network excitability. Yet the specific contributions of astrocytic gap-junction coupling and potassium spatial buffering to neuronal synchronization across different spatial scales remain poorly understood. To address this gap, we developed a microcircuit biophysical model consisting of two astrocyte-neuron modules, each comprising one astrocyte coupled to five neurons. Astrocyte-neuron interactions are mediated exclusively through shared extracellular potassium dynamics. Using a reduced astrocyte model that captures both local membrane and syncytial potassium buffering, we systematically investigated how astrocytic potassium handling shapes neuronal activity patterns and inter-module synchronization. Our results demonstrate that astrocytes prevent the emergence of pathological states — such as sustained ictal activity and depolarization block, by stabilizing extracellular potassium levels. Furthermore, we show that astrocytic gap-junction coupling strength critically regulates phase synchronization between neuronal modules: stronger coupling promotes inter-module synchrony under physiological conditions, whereas impaired astrocytic function drives networks toward pathological hypersynchronization when extracellular potassium is elevated. These findings support the hypothesis that astrocytic networks impose modularity on hippocampal neuronal assemblies, and suggest that astrocytic connexins may represent a relevant therapeutic target in epilepsy and other disorders characterized by aberrant neural synchronization.

**Author summary**

## Introduction

Neural synchronization has been proposed as a communication mechanism in both the resting-state and in task-related conditions [1]. Synchronization plays an important role in normal aging [2] but excessive synchronization can be deleterious, for example in epilepsy [3]. Among epileptic disorders, temporal lobe epilepsy is the most common form in adults, with the hippocampus, and especially the mesial temporal structures representing one of the most epileptogenic regions [4]. In epilepsy, the role of extracellular potassium ([*K*^+^]_*o*_) concentration is central for the onset and the continuation of epileptic activities, as demonstrated by numerous experimental and theoretical studies (See [5] for a review). In particular, it is well known that hippocampal slices subjected to low external calcium concentration, which blocks synaptic transmission, can sustain non-synaptic epileptiform activity [6–9]. In this experimental model, the removal of chemical synapses unmasks non-synaptic synchronization mechanisms in which elevated extracellular *K*^+^ accumulation plays a critical role [10]. Non-synaptic epileptiform activity is a term that encompasses all the mechanisms of seizure generation and propagation that are independent of chemical synapses, including electrical field effects (i.e., ephaptic interactions), gap junctions, and ionic diffusion through the extracellular space. The relative contribution of these mechanisms, as well as the processes governing initiation, propagation, and termination of non-synaptic ictal events, remain topics of ongoing investigation [11]. Multiple in vitro and in vivo studies demonstrate that the extracellular space dynamics play a central role in non-synaptic ictal initiation and propagation [12–14]. The hippocampus is particularly susceptible to such mechanisms due to its dense packing of neuronal cell bodies and highly parallel cellular architecture: the same structural features that also underlie the characteristically large field potentials recorded in this region [11, 15–17]. Among the ionic mechanisms contributing to non-synaptic epileptogenesis, extracellular *K*^+^ accumulation plays a particularly important role in generating and sustaining synchronized ictal activity in the hippocampus [18, 19], as corroborated by computational modelling studies [20–22]. Given the critical role of *K*^+^ dynamics, recent experimental evidence suggests that non-optimal astrocytic *K*^+^ buffering mechanisms may be one of the main drivers of non-synaptic ictal activity in the hippocampus [23, 24].

Accumulating evidence demonstrates the role of astrocytes in modulating the synchronizability of neuronal populations (see [25] for a review). However, the conditions under which potassium buffering effectively prevents seizure activities remain incompletely understood. Effective regulation of extracellular *K*^+^ depends on three complementary astrocytic mechanisms: inward-rectifying *Kir*4.1 channels, the *Na*^+^*/K*^+^-ATPase pump, and the connexin-mediated gap-junction coupling [26–31]. Together, these mechanisms enable the astrocytic syncytium to redistribute excess extracellular potassium, thereby maintaining ionic homeostasis and near-isopotentiality across the astrocytic network [32]. Disruption of any of these components promotes network hyperexcitability and pathological hypersynchronization, increasing seizure susceptibility [30, 33–35]. However, two fundamental questions remain unresolved: how is *Kir*4.1 expression dynamically regulated, and what degree of gap-junction coupling optimally balances efficient *K*^+^ clearance without inducing excessive neuronal synchronization, particularly in epileptogenic contexts [36–39] ?

In the hippocampus specifically, astrocytes form extensive, spatially organized syncytial networks that modulate local circuit dynamics and information processing [40], suggesting that they may (dynamically) shape the modular organization of neuronal assemblies [40–42]. Motivated by these findings, we hypothesize that astrocytes can impose and regulate functional modularity within small neuronal assemblies designed to mimic the dense cellular packing characteristic of hippocampal tissue.

To investigate how astrocytic potassium buffering shapes neural synchronization and modulates network activity under elevated extracellular potassium concentrations, a hallmark of epileptogenic conditions, we developed a biophysical hippocampal microcircuit model incorporating explicit potassium dynamics in both neurons and astrocytes. The neuronal component employs a reduced Hodgkin-Huxley model [43] with explicit potassium dynamics proven to trigger the full spectrum of potassium-driven activity states [44], while the astrocytic component is based on a recently developed model capable of capturing potassium buffering dynamics under diverse pharmacological conditions [45]. The microcircuit scale bridges the mechanistic gap between single-cell and mesoscale descriptions of brain dynamics, and the compact cytoarchitecture of the hippocampus makes it an ideal substrate for examining how local astrocyte-neuron ensembles respond to extracellular potassium fluctuations at fine spatial scales [16, 17]. We first characterize the dynamical repertoire of the network with and without astrocytic modulation, then systematically quantify how astrocytic gap junction coupling and potassium buffering regulate the emergence and spatial organization of neural synchronization across physiological and pathological regimes.

## Results

In the first part of this section, we introduce the overall architecture of the model, with particular emphasis on parameters that form the focus of our analysis. We then characterize the dynamical repertoire of the network under two conditions: with and without astrocytic modulation. Finally, we quantify the contribution of astrocytes to network-level synchronization and assess how their activity shapes the emergence of synchronized neural dynamics.

### Astrocyte-Neurons Model

We implemented a microcircuit model comprising two coupled astrocyte-neuron modules. Within each module *M*, with *M* denoting *A* or *B*, the astrocyte interacts with neurons through a shared extracellular potassium pool [*K*^+^]*o*^*M*^ in the extracellular space (ECS) compartment, where the subscript “o” denotes extracellular (outside). The model architecture is illustrated in Fig. 1. To reflect hippocampal anatomy, each module incorporates an astrocyte-to-neuron ratio of 1 : 5, consistent with experimental observations [46]. The extracellular potassium concentration in module *M* (Eq. 40) integrates contributions from five distinct sources: (i) the baseline concentration 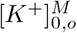 at *t* = 0, (ii) the astrocytic potassium flux 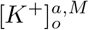, (iii) the neuronal potassium release summed across all five neurons 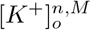, (iv) the exchange with the external bath (EB) 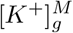, and (v) the diffusive coupling between modules 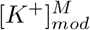. The governing equations and parameters for each component are detailed in the *Methods* section. The two modules interact through three pathways: (1) synaptic connections between neurons, (2) potassium diffusion within the two ECS compartments, and (3) a gap-junction coupling the two astrocytes. Neurons within and across modules are arranged in a fully connected network, a choice justified by the small size of the network (10 neurons). To increase biological plausibility, an inhibitory-excitatory ratio of 1 : 4 is chosen. Excitatory synaptic conductances are taken from a uniform distribution between [0.5, 0.7] *nS*, whereas inhibitory conductances were drawn from a uniform distribution between [1, 3] *nS*. These ranges were selected based on in-silico calibrations using hippocampal experimental data [47]. The ECS compartments of the two modules were coupled via a potassium flux governed by a time constant (*ϵ*_*mod*_ in Eq. (39)) that is ten times less than the constant that governs the potassium flux between ECS and EB (*ϵ* in Eq. (34)). This choice reflects experimental evidence that diffusion between distinct cellular layers is significantly impeded by anatomical and molecular barriers. Brain extracellular space tortuosity arises from geometric constraints imposed by densely packed cells, the presence of extracellular matrix components that physically obstruct ion passage, and local variations in ECS architecture, including dead-space microdomains [48–50]. Together, these factors substantially reduce the effective diffusion coefficient for ions moving between neuronal assemblies compared to exchange with the well-mixed external bath. Astrocytes communicate to each other through a gap-junction represented schematically in Fig. 1. The gap-junction conductance (*g*_*gap*_) was the primary parameter varied to assess how astrocytic coupling influences neural synchronization.

**Fig 1.**
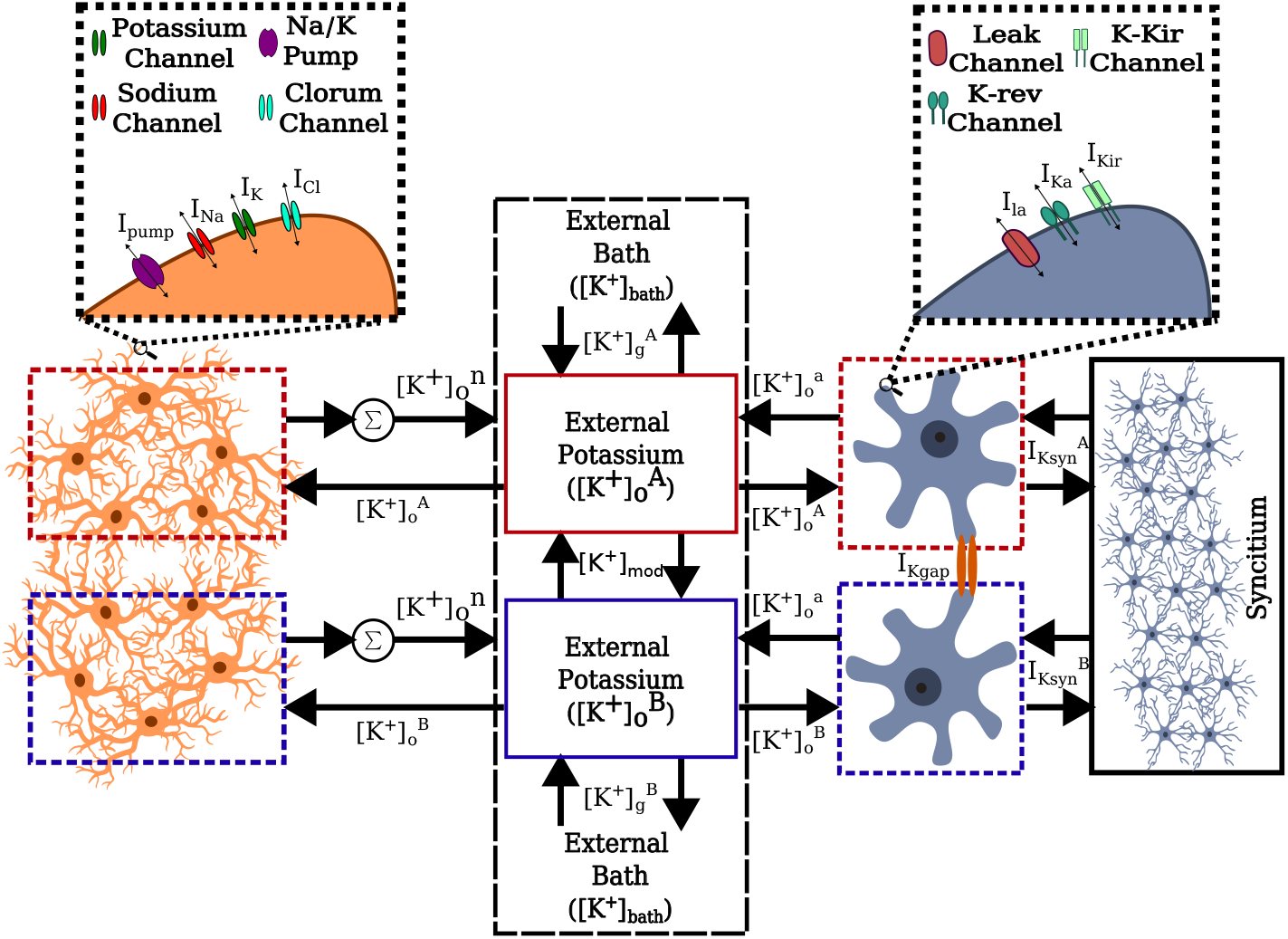
Model scheme: red squares/rectangles indicate module *A*, blue squares/rectangles indicate module *B*, and black squares/rectangles represent compartments common to both modules.

Finally, to break the symmetry between modules and prevent artificially synchronized initial states, all state variables were initialized by random sampling from physiologically plausible ranges, as described in the *Methods* section. Additional mathematical details and further experimental choices are also provided in the *Methods* section.

### Dynamical Repertoire

First, we systematically investigated how astrocytic potassium buffering shapes the dynamical repertoire of neuronal membrane potentials. This analysis established a baseline description of the network’s intrinsic activity and enabled us to evaluate how these activity patterns evolve across different extracellular potassium conditions. The resulting behaviors, and in particular the role of astrocytes in shaping them, are presented in Figure 2.

**Fig 2.**
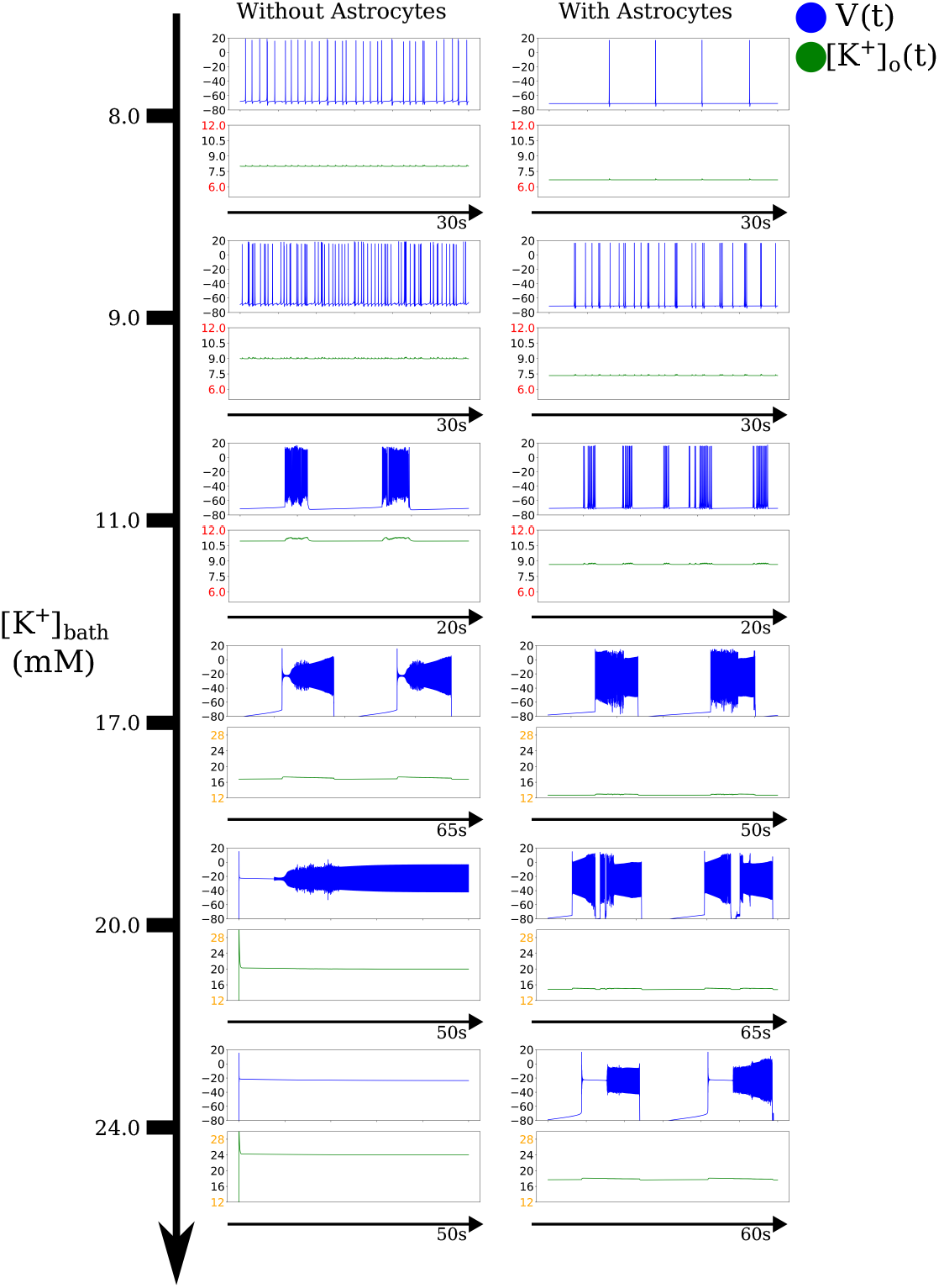
Dynamical repertoire of the first neuron in module *B* with and without astrocytes. The plots show the membrane potential (blue traces, in *mV*), and the extracellular potassium concentration (green traces, in *mM*) as [*K*^+^]_*bath*_ varies. The cases without astrocytes were previously identified in [44, 51]. Due to the large differences observed in the amplitudes of [*K*^+^]_*o*_, two different y-axis ranges were selected: the first spanning [6, 12] *mM*, and the second [12, 28] *mM*, highlighted in red and orange, respectively.

In the absence of astrocytes, progressively increasing [*K*^+^]_*bath*_ allowed the neuronal network to reproduce the full spectrum of activity regimes previously identified in the single-neuron model [44], and in a large neural network model [51]. As [*K*^+^]_*bath*_ increases, the network passes from stable resting states to spike train, tonic spiking, bursting, Seizure-Like Event (SLE), Sustained Ictal Activity (SIA), and, eventually, Depolarization Block (DB). These transitions occurred in a graded manner and were accompanied by marked elevations in extracellular potassium concentration, consistent with classical dynamical signatures of potassium-driven excitability.

The presence of astrocytes, however, profoundly altered this progression. When astrocytic dynamics were included, increasing [*K*^+^]_*bath*_ no longer drove the network into SIA or DB. Instead, before the system could reach these extreme activity states, astrocytic potassium buffering mechanisms became strongly activated. This buffering effectively stabilized extracellular potassium levels and prevented the runaway accumulation necessary for the emergence of highly depolarized or pathological regimes. As a result, the network remained confined to lower-excitability states, demonstrating the robust homeostatic influence of the astrocytic component.

These differences are clearly illustrated in Fig. 2, which compares the membrane potential of the first neuron and the corresponding extracellular potassium concentration across a range of [*K*^+^]_*bath*_ values in both astrocyte-free and astrocyte-present conditions. Simulations were run for a total duration of *t*_*max*_ = 120 *s* with an integration time step of *T* = 0.03 *ms*.

### Astrocytes as Synchronizers of Neural Activity

Since noise is absent in the model, the network exhibits elevated baseline synchrony despite neurons being initialized with different initial conditions. To increase asynchrony between the activity of neurons in the two modules and isolate the effect of the astrocytic gap-junction on neural synchronization, we set the syncytium coupling parameter asymmetrically: 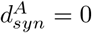 in module *A* and 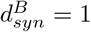 in module *B*. This configuration results in one astrocyte being connected to the syncytium while the other remains isolated, allowing us to assess the specific contribution of inter-astrocytic gap-junction strength (*g*_*gap*_) to network synchronization.

To investigate the effect of astrocytic coupling in the neural network dynamics, we focus on the two main parameters that regulate the potassium buffering within the astrocyte, namely (1) the threshold of the sigmoid function that controls potassium exchange between an astrocyte and the syncytium, as well as with the neighbouring astrocyte (*Z*_*th*_ in Eq. (31)); and (2) the gap-junction conductance *g*_*gap*_ between the two astrocytes (see Eq. 27).

To quantify the relative strength of synchronization within versus between modules, we defined the within/between Phase Locking Value (PLV) ratio (*WB*_*PLV*_), defined as the average intra-module PLV divided by the inter-module PLV. Specifically, we first calculated the mean PLV among all neuron pairs within module *A* (⟨*PLV*⟩_*A,A*_) and within module *B* (⟨*PLV*⟩_*B,B*_), then averaged these two values. We then divided this average intra-module synchrony by the mean PLV computed between all neuron pairs across the two modules (⟨*PLV*⟩_*A,B*_). Higher *WB*_*PLV*_ values indicate that neurons are more synchronized within their respective modules than between modules, whereas lower values indicate that inter-module synchronization approaches intra-module levels. Detailed methodology is provided in the *Methods* section.

The results of this study are summarized in Fig. 3. Panel (a) illustrates how *WB*_*PLV*_ varies as a function of *g*_*gap*_ and the *Z*_*th*_ across different extracellular potassium concentrations ([*K*^+^]_*bath*_). As [*K*^+^]_*bath*_ increases, the system transitions through distinct dynamical regimes (Fig. 2), revealing that the role of astrocytic coupling in modulating inter-module synchronization depends critically on both network excitability and the functional status of astrocytic potassium buffering.

**Fig 3.**
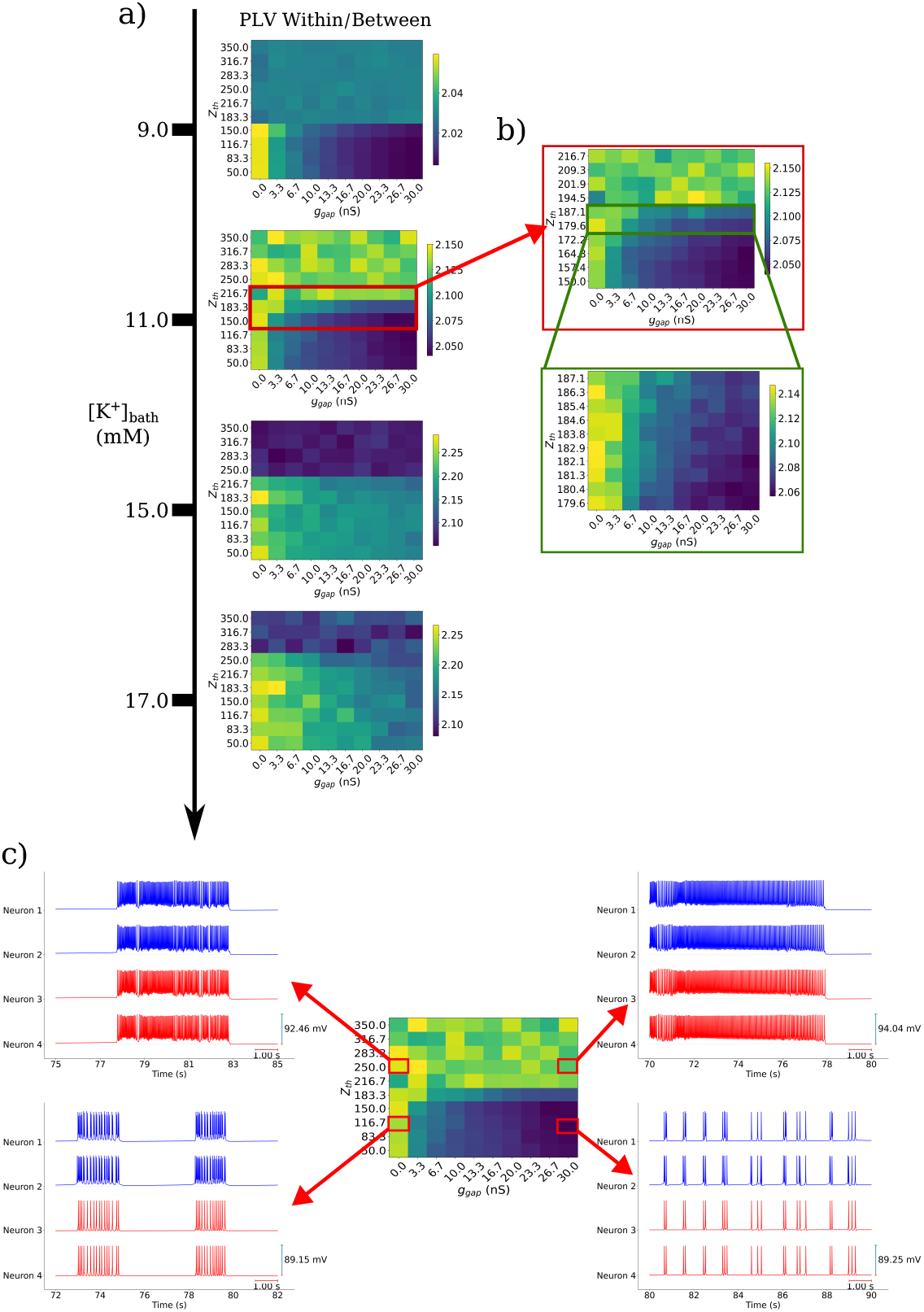
Astrocytes as phase synchronizers of neural activity. **(a)** PLV within/between (*WB*_*PLV*_) as a function of *Z*_*th*_ (threshold of the sigmoid function regulating potassium exchange with the astrocytic syncytium, unitless) and *g*_*gap*_ (astrocytic gap-junction conductance, in *nS*), across different dynamical regimes obtained by varying [*K*^+^]_*bath*_. **(b)** Transition of *WB*_*PLV*_ with increasing *Z*_*th*_ for a fixed [*K*^+^]_*bath*_ = 11 *mM* . **(c)** Neuronal synchronization states corresponding to distinct *WB*_*PLV*_ values (i.e., different combination of *Z*_*th*_ and *g*_*gap*_) showing the membrane potentials of neurons in module *A* (blue) and module *B* (red) for [*K*^+^]_*bath*_ = 11 *mM* .

In the spike train regime ([*K*^+^]_*bath*_ = 9 *mM* ), neurons exhibit regular firing patterns with relatively high baseline synchronization even without astrocytic gap-junction coupling. Under these conditions, increasing *g*_*gap*_ produces only a modest enhancement of inter-module synchronization, suggesting that synaptic interactions dominate network coordination in physiological regimes. However, as [*K*^+^]_*bath*_ increases to intermediate values ([*K*^+^]_*bath*_ = 11 *mM* ), corresponding to bursting activity, the influence of the astrocytic coupling becomes substantially more pronounced. Here, increasing *g*_*gap*_ drives a marked transition from modular organization (high *WB*_*PLV*_ ) toward global network synchronization (low *WB*_*PLV*_ ), demonstrating that astrocytes can actively reshape the functional architecture of neuronal ensembles through potassium spatial buffering. This effect becomes pronounced the most under pathological conditions ([*K*^+^]_*bath*_ ≳ 15 *mM* ). In this state, astrocytic gap-junction coupling acts as a powerful synchronization mechanism: networks with weak or absent coupling (*g*_*gap*_ ≈ 0 *nS*) maintain relative independence between modules, whereas strong coupling (*g*_*gap*_ ≈ 30 *nS*) drives the entire network into highly synchronous activities. However, as *Z*_*th*_ increases, potassium exchange between the astrocyte and the syncytium, as well as between astrocytes across modules, becomes progressively impaired. Higher *Z*_*th*_ decreases synchronization under physiological conditions, while still permitting the emergence of strongly synchronized pathological states in the presence of elevated extracellular potassium. These findings support the hypothesis that astrocytic dysfunction contributes to the maintenance of non-synaptic hippocampal activity during seizures.

Panel (b) examines the *Z*_*th*_-dependent transition in more detail for a fixed extracellular potassium concentration ([*K*^+^]_*bath*_ = 11 *mM* ). This analysis reveals that the shift from optimal astrocytic performance (low *Z*_*th*_) to functional impairment (high *Z*_*th*_) is gradual rather than abrupt, with intermediate *Z*_*th*_ values producing graded changes in synchronization patterns.

Finally, panel (c) provides exemplar neuronal voltage traces at different *WB*_*PLV*_ levels, illustrating the mechanistic link between astrocytic coupling and network dynamics for a fixed extracellular potassium concentration ([*K*^+^]_*bath*_ = 11 *mM*). At high *WB*_*PLV*_ values (dysfunctional astrocytes with low *g*_*gap*_ or high *Z*_*th*_), neurons in the two modules show asynchronous bursting activity, with a longer single burst duration when *Z*_*th*_ is high. As *g*_*gap*_ increases and *WB*_*PLV*_ decreases (for *Z*_*th*_ ≲ 216.7), neurons across modules fire in phase-locked patterns with regular spiking activity. Notably, when astrocytes function correctly, network activity shifts toward dynamical regimes characteristic of lower [*K*^+^]_*bath*_, suggesting that astrocytic coupling influences both temporal coordination and the intensity and persistence of network activities. Taken together, these results reveal two functionally distinct, yet mechanistically coupled, roles of astrocytic potassium buffering in shaping network dynamics. First, astrocytes act as homeostatic regulators, preventing the runaway extracellular potassium accumulation that drives transitions toward sustained ictal activity and depolarization block in the absence of glial buffering. Second, and beyond this protective function, astrocytic gap-junction coupling emerges as an active determinant of the spatial architecture of synchronization: by redistributing excess extracellular potassium across modules, the astrocytic syncytium modulates the balance between intra- and inter-module phase locking in an excitability-dependent manner. Critically, these two roles are not independent: the same connexin-mediated coupling that promotes inter-module synchronization under physiological conditions becomes a driver of pathological hypersynchronization when extracellular potassium is chronically elevated and astrocytic buffering capacity is compromised. These aspect are modeled by a sigmoid threshold *Z*_*th*_, which governs potassium exchange between the astrocyte and the syncytium, mediating a continuous (rather than abrupt) transition between these regimes. In other words, the evidence provided suggests that astrocytic dysfunction need not be complete to substantially alter network coordination. Collectively, these findings indicate that the astrocytic syncytium does not merely respond passively to neuronal activity but might actively coordinate neuronal assemblies into functional modules whose degree of coupling is dynamically regulated by the state of extracellular potassium homeostasis. As such, this mechanism would have direct implications for the transition between physiological synchronization and pathological patterns, such as seizure-like states.

## Discussion

To investigate how astrocytic potassium buffering shapes local neural synchronization, we developed a hippocampal microcircuit model and analyzed its impact on network activity. In this framework, each astrocyte modulates the activity of multiple neurons through a shared extracellular potassium pool (Fig. 1), consistent with the anatomical organization of astrocyte-neuron interactions across brain regions [46, 52–54]. We also took into account the extracellular tortuosity between modules by setting the inter-module diffusion flux as one order of magnitude smaller than the exchange flux between each module and the external bath. This is consistent with experimental measurements showing that dense cellular packing substantially impedes ion diffusion between neuronal assemblies [48–50].

This architecture allows us to investigate how glial cells shape local neuronal interactions and how these microscale effects translate into coordinated population activity, a transition whose underlying mechanisms remain unclear. We hypothesize that glial modulation of extracellular potassium buffering critically regulates neuronal communication and coordination. Our results demonstrate that astrocytic potassium buffering exerts two distinct yet complementary influences on network dynamics: a homeostatic role that prevents pathological activity, and an organizational role that regulates the spatial structure of synchronization.

In the absence of astrocytes, we found that the progressive increase in the external bath potassium concentration raise extracellular potassium and drive the network through the entire spectrum of pathological states, from seizure-like events to sustained ictal activity and depolarization block, consistent with experimental observations [18, 19] and previous computational models [21, 22, 44]. However, when astrocytic dynamics were included, the network remained confined to physiological or mildly hyperexcitable regimes, demonstrating that astrocytic potassium clearance mechanisms effectively stabilize network activity (Fig. 2).

Beyond this homeostatic function, our findings reveal that astrocytic gap-junction coupling crucially regulates inter-module synchronization in an excitability-dependent manner. In physiological regimes with low network firing rates (regular spiking) synaptic interactions dominate, and astrocytic coupling produces modest effects. As extracellular potassium increases to intermediate levels ([*K*^+^]_*bath*_ = 11 *mM*), astrocytic coupling becomes a powerful regulator, driving transitions from modular organization to global synchronization (Panel a) Fig. 3). This extends previous work showing that astrocytes modulate firing patterns [23, 24] by demonstrating that they actively shape the spatial architecture of network synchronization.

Furthermore, the sigmoid threshold *Z*_*th*_, which regulates potassium exchange between astrocytes and the syncytium, mediates a qualitative shift in network behaviour (Panels b) and c) Fig. 3). Under physiological conditions, elevated *Z*_*th*_ (mimicking impaired connexin function) maintains modular organization, whereas under elevated potassium, impaired astrocytic function facilitates strongly synchronized pathological states. Such dual role suggests that astrocytic dysfunction contributes differentially to seizure susceptibility depending on baseline excitability, consistent with experimental observations linking astrocytic pathology to epileptogenesis [30, 33, 35]. These findings support the hypothesis that astrocytic networks impose modularity on hippocampal neuronal assemblies [40, 41] and suggest that targeting astrocytic connexins may offer cell-type-specific therapeutic strategies for epilepsy and other disorders characterized by aberrant synchronization [55].

Although our model incorporates several features that enhance its biological plausibility, including astrocytes influencing multiple neurons and reduced diffusion between the extracellular spaces of the two modules due to tortuosity, several important limitations must be acknowledged. The model includes only ten fully connected neurons and focuses exclusively on potassium dynamics, omitting mechanisms such as glutamate uptake, calcium signalling, activity-dependent plasticity, and astrocytic heterogeneity, all or which may interact synergistically (see [56–61] for comprehensive reviews of astrocytic computational models). Additionally, the two-module architecture limits investigation of multi-scale coordination, and the absence of explicit stochastic elements prevents exploration of how astrocytic networks filter or amplify noise in neural circuits.

A natural extension is bridging microcircuit and larger-scale descriptions of brain dynamics. Developing a mesoscale model by coupling multiple modules arranged to reflect hippocampal laminar organization could investigate how astrocytic buffering regulates ictal propagation across layers and how layer-specific dysfunction contributes to seizure generation. Another promising direction involves deriving a neural mass model that captures essential astrocyte-mediated potassium buffering dynamics while reducing computational complexity. Recent biophysically inspired mean-field models that incorporate ion exchange mechanisms [51] provide a foundation for this approach. Extending this framework to include spatially distributed astrocytic gap-junction coupling could enable simulation of how astrocytic networks shape large-scale brain dynamics and epileptiform activity propagation across cortical regions [60].

In conclusion, this work demonstrates that astrocytic gap-junctions critically regulate neural synchronization, supporting the view that astrocytic networks might represent legitimate therapeutic targets for treating disorders characterized by aberrant neural synchronization, such as epilepsy. Our microcircuit model provides a quantitative framework for understanding how local astrocyte-neuron interactions scale to network-level phenomena and establishes groundwork for investigating how astrocytic networks coordinate dynamics from microcircuits to whole-brain activity.

## Methods

In this section, we present the mathematical formulation of the proposed model along with a detailed description of the analysis methods employed. The aim is to provide a comprehensive account of the underlying equations, parameters, and computational procedures used to investigate the neural–astrocytic dynamics and their synchronization properties.

### Model

The used model is composed of two modules, and it is represented in Fig. 1. Each module contains three main parts: i) a neuronal network, ii) one astrocyte, embedded into iii) the extracellular space.

The extracellular space (ECS) is described by the dynamics of the extracellular potassium, which links the activity of neurons with that of astrocytes and is directly related to the extracellular bath. Each part of the model is described in detail in the following subsections.

### Neural Model

The single-neuron model is a Hodgkin–Huxley (HH)-like model taken from the work of Depannemaecker et al. [44]. It is a three-compartment model, with the three compartments indicating: Intracellular Space (ICS), Extracellular Space (ECS), and, External Bath (EB).

The ICS and ECS compartments communicate through: currents flowing through *Na*^+^, *K*^+^, and *Cl*^−^ voltage-gated channels (*I*_*Na*_, *I*_*K*_, and *I*_*Cl*_) and the sodium-potassium pump generated current (*I*_*pump*_).

The original model is a slow-fast dynamical system based on 4 differential equations associated with membrane potential, gating variable, rate of change of the intracellular potassium and diffusion of potassium from the bath. Here we considered the 3 first differential equations for the neuronal dynamics, the fourth being adapted at module level. The fast variables are the membrane potential (*V*) and the potassium conductance gating variable (*n*). While the slow variable is the intracellular potassium concentration variation (Δ[*K*^+^]_*i*_), where the subscript “i” stands for *inside*, i.e., intracellular. The model equations for the ICS compartment are:

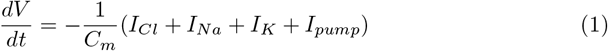

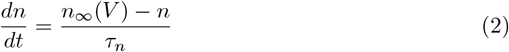

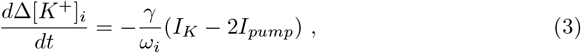

being *γ* the conversion factor. This is a scaling parameter that permits the inclusion of all the mechanisms, not detailed in this model, affecting the potassium concentration variation (such as co-transporters, exchangers, etc). It has the same unit as the inverse Faraday constant (*mol/C*). Parameters, physiological reference, and initial values are reported in Table 2, 3, and 4, respectively.

**Table 1.**
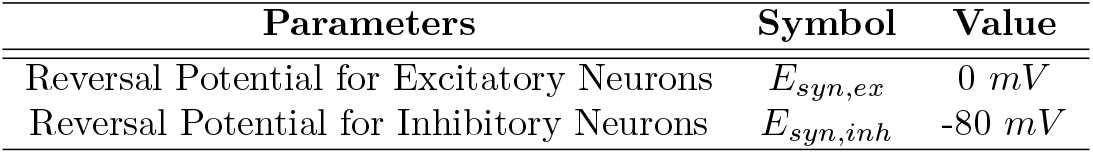
Synaptic Parameters Values.

**Table 2.** Parameters Values.

| Parameters | Symbol | Value |
| --- | --- | --- |
| *Neuron Parameters* |  |  |
| Membrane Capacitance | $C_m$ | 40 nF |
| Gating Time Constant | $\tau_n$ | 10 ms |
| Chloride Conductance | $g_{Cl}$ | 7.5 nS |
| Maximal Potassium Conductance | $g_K$ | 22 nS |
| Maximal Sodium Conductance | $g_{Na}$ | 40 nS |
| Potassium Leak Conductance | $g_{K,l}$ | 0.12 nS |
| Sodium Leak Conductance | $g_{Na,l}$ | 0.02 nS |
| Intracellular Volume | $\omega_i$ | 2160 $\mu m^3$ |
| Extracellular Volume | $\omega_o$ | 720 $\mu m^3$ |
| Intra/Extra-Cellular Volume Ratio | $\beta = \omega_i/\omega_o$ | 3 |
| Conversion Factor | $\gamma$ | 0.002 mol/C |
| Maximal Na/K Pump Current | $\rho$ | 250 pA |
| *Astrocyte Parameters* |  |  |
| Membrane Capacitance | $C_{m,a}$ | 100 nF |
| Maximal potassium conductance of Kir channel | $g_{Kir}$ | 10 nS |
| Pseudo parameter | $A$ | 1 mM$^{-\frac{1}{2}}$ |
| Potassium Astrocytes Conductance | $g_{K_a}$ | 0 nS |
| Potassium Leak Conductance Astrocytes | $g_{l_a}$ | 0.1 nS |
| Potassium Conductance between $m$ and $n$ Astrocytes | $g_{gap}$ | 0.0-30. nS |
| Surface | $\sigma_a$ | 1600 $\mu m^2$ |
| Intracellular Astrocyte Volume | $\omega_a$ | 2000 $\mu m^3$ |
| Conversion Factor between External/Internal | $\gamma_t$ | 5 m$^{-2}$ |
| Conversion Factor between Syncytium/Internal | $\gamma_s$ | 6 m$^{-2}$ |
| Membrane Potential Syncytium | $V_{a,s}$ | -90 mV |
| Reversal Leak Potential | $E_{l_a}$ | -70 mV |
| Membrane potassium permeability | $P_K$ | 6e$^{-5}$ m$^3/s$ |
| gap-junction with Syncytium coupling strength | $d_{syn}$ | 1 |
| Sigmoid Parameter | $Z_{th}$ | 50-350 |
| Sigmoid Parameter | $Z_s$ | 0.05 |
| *ECS Parameters* |  |  |
| Diffusion Time Constant with EB | $\epsilon$ | 0.01 ms$^{-1}$ |
| Diffusion Time Constant Between Modules | $\epsilon_{mod}$ | 0.001 ms$^{-1}$ |

**Table 3.** Physiological Reference Values.

|  | Ion | Concentration |
| --- | --- | --- |
| *Neuron Reference Values* |  |  |
| **Extracellular** | $[Na^+]_{0,o}$ | 138 mM |
| | $[Cl^-]_{0,o}$ | 112 mM |
| **Intracellular** | $[K^+]_{0,i}$ | 140 mM |
| | $[Na^+]_{0,i}$ | 16 mM |
| | $[Cl^-]_{0,i}$ | 5 mM |
| *Astrocyte Reference Values* |  |  |
| **Intracellular** | $[K^+]_{0,a}$ | 135 mM |
| *ECS Reference Values* |  |  |
| **External Bath** | $[K^+]_{bath}$ | [7-25] mM |
| **Extracellular** | $[K^+]_{0,o}$ | 4.8 mM |

**Table 4.** Initial Values.

| Variable | Initial Value |
| --- | --- |
| *Neuron Initial Values* |  |
| $[Na^+]_o$ | $[Na^+]_{0,o}$ |
| $[Cl^+]_o$ | $[Cl^-]_{0,o}$ |
| $[K^+]_i$ | $[K^+]_{0,i}$ |
| $[Na^+]_i$ | $[Na^+]_{0,i}$ |
| $[Cl^-]_i$ | $[Cl^-]_{0,i}$ |
| $\Delta[K^+]_i$ | [-3, 0.2] mM |
| $[K^+]_g$ | [-0.3, 2.] mM |
| $V$ | [-85, -65] mV |
| $n$ | $n_\infty(V_0)$ |
| *Astrocyte Initial Values* |  |
| $[K^+]_a$ | $[K^+]_{0,a}$ |
| $V_a$ | [-95, -75] mV |
| $[K^+]_{a,t}$ | [-3, 0.5] mM |
| $[K^+]_{a,s}$ | [-2, 2] mM |
| *ECS Initial Values* |  |
| $[K^+]_o$ | $[K^+]_{0,o}$ |
| $[K^+]_g$ | [-0.3, 2.] mM |
| $[K^+]_{o,mod}$ | 0 mM |

The current contributions at the right-hand side of Eq. (1) are given by the following expressions:

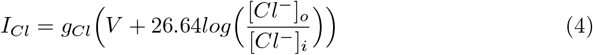

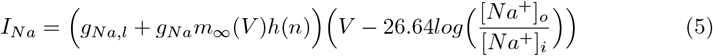

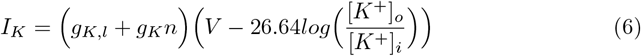

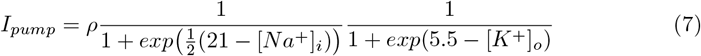

The conductance variables depend on the membrane potential *V* and the gating variable *n*:

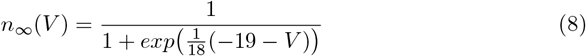

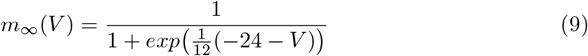

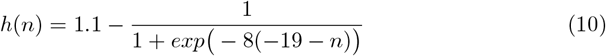

The concentration variation of potassium and sodium can be actualised by assuming a constant chloride concentration and local equilibrium:

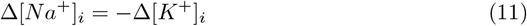

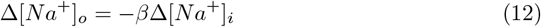

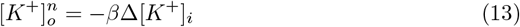

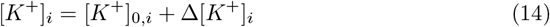

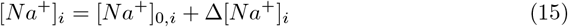

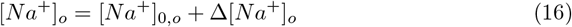

To link neurons in a neural network, a synaptic current term is added to Eq. (1), which is written as a conductance-based model. Then, considering the *j*-th neuron, we get:

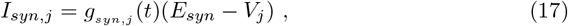

being *g*_*syn,j*_ the synaptic of the *j*-th neuron, *V*_*j*_ the postsynaptic potential of the *j*-th neuron, and *E*_*syn*_ the synaptic reversal potential. Typical values of *E*_*syn*_ are reported in Table 1. As in [62], this last equation can be written as:

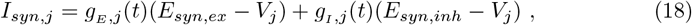

being *g*_*E*_ (*t*) and *g*_*I*_ (*t*) the excitatory (E) and inhibitory (I) conductances, respectively, which increase by quantities *Q*_*E*_ = [0.5 − 0.7] *nS* and *Q*_*I*_ = [1 − 3] *nS* for each incoming spike. These values are set according to the physiological values found in the hippocampus [47]. The increment of E/I conductance is followed by an exponential decrease according to the following linear relaxation equations:

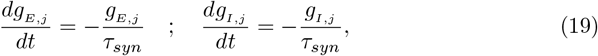

where *τ*_*syn*_ = 5 *ms*.

### Astrocytic Model

The astrocyte model [45] is composed of three differential equations that represent the variation through time of: the astrocyte Membrane Potential (*V*_*a*_) (eq.:20, the potassium concentration exchanged through the membrane ([*K*^+^]_*a,t*_) (eq: 21, the potassium concentration exchanged with the syncytium ([*K*^+^]_*a,s*_) (eq: 22). The equations for the *k*-th astrocyte are:

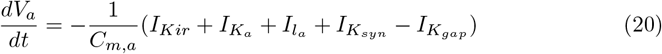

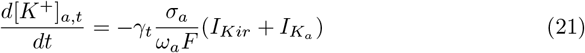

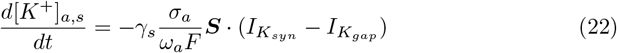

Where the current equations are:

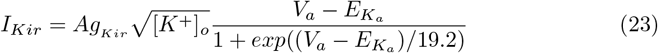

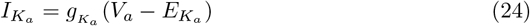

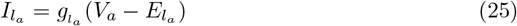

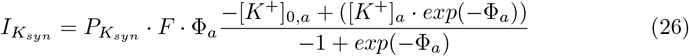

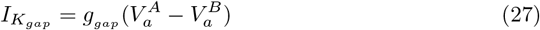

Where 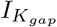 is the current that comes from the astrocyte *n* to the astrocyte *m* through a gap-junction. Moreover:

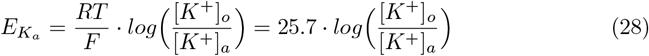

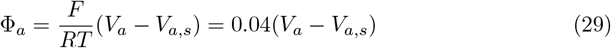

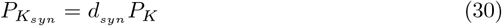

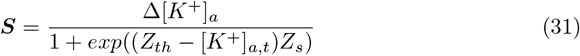

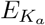 is the potassium Nernst potential of the *m*-th astrocyte, Φ_*a*_ the potential difference between the *m*-th astrocyte and the syncytium, 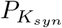 the average cell permeability, and ***S*** the sigmoid function regulating the opening of the gap-junction between syncytium/other astrocytes and the *k*-th astrocyte. The potassium concentration variation for the *k*-th astrocyte is calculated as follows:

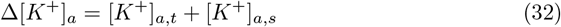

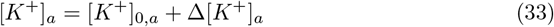

Parameters, physiological reference and initial values are reported in Table 2, 3, and 4, respectively.

### Extracellular Space

The ECS of the *M*-th module is linked to the EB through a differential equation that describes the extracellular potassium buffering by the EB 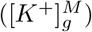:

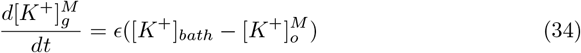

Being [*K*^+^]_*bath*_ the potassium concentration in the EB, and 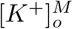 the extracellular potassium concentration of the *M*-th module. Within each module, astrocytes and neurons are coupled through their shared extracellular potassium pool. The combined astrocytic and neuronal contribution to 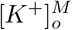 is:

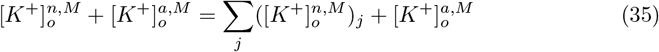

Where 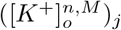 denotes the extracellular potassium contribution from neuron *j* in module *M*, and 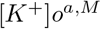 denotes the astrocytic contribution in module *M*. These are defined as:

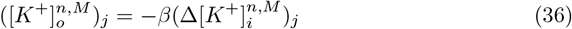

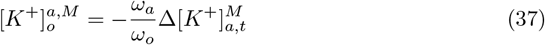

Finally, the extracellular potassium concentration in module *M* is given by the sum of all contributions:

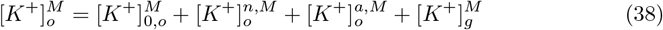

Where 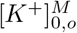 is the initial extracellular potassium concentration at *t* = 0. When multiple modules are present, inter-module diffusion must be included. The term [*K*^+^]*mod*^*M*^ represents potassium exchange between modules *M* and *N*:

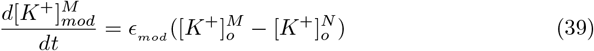

*ϵ*_*mod*_ is a constant parameter that regulates the exchange of potassium between the two modules. Then, Eq. (38) finally became:

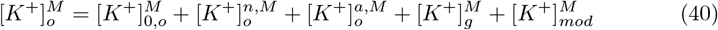

Parameters, physiological reference, and initial values are reported in Table 2, 3, and 4, respectively.

### Measure of Neural Synchrony: Phase Locking Value

The Phase Locking Value (PLV) is a widely used metric for quantifying phase synchronization in nonlinear dynamical systems. Since the brain exemplifies nonlinear dynamical systems, the PLV provides an effective means of assessing interactions among neural populations or brain regions, as observed in Local Field Potentials (LFPs), EEG and MEG [63–65].

There are various methods for extracting phases from a continuous signal. In this work, the most common technique based on the analytic signal was applied [66]. Starting from the neural voltage signal of the *i*-th neuron (*V*_*i*_(*t*)), the analytic signal is defined as:

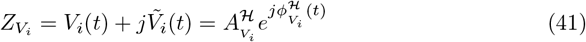

where 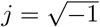, and 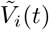 is the Hilbert Transform of *V*_*i*_(*t*):

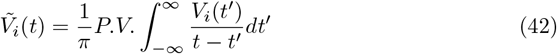

Being *P*.*V*. the Cauchy principal value. From *Z* the Hilbert phase can be extracted as:

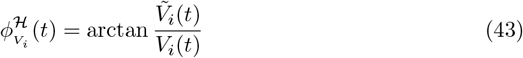

In the same way the Hilbert phase for the *k*-th neuronal voltage can be extracted 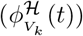. The average *PLV* between a couple of neuronal voltages *V*_*i*_ and *V*_*k*_ is then defined as:

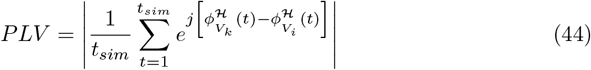

By doing this for each couple of neurons in the network the *PLV* matrix was developed. To assess the level of synchronization between modules, and to compare it with the synchronization within each module, we evaluate the within/between Phase Locking Value (PLV), defined as:

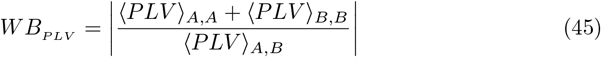

Here, ⟨*PLV* ⟩_*A,A*_ (and analogously ⟨*PLV* ⟩_*B,B*_) denotes the mean *PLV* computed among neurons within module *A* (or *B*), whereas ⟨*PLV*⟩ _*A,B*_ represents the mean *PLV* computed between neurons belonging to the two different modules.

## Acknowledgments

This work was supported by the Next-Generation-EU programme under the funding schemes PNRR-PE-AI scheme (M4C2, investment 1.3, line on AI) FAIR “Future Artificial Intelligence Research”, grant id PE00000013, Spoke-8: Pervasive AI. This research also received funding from the European Union’s Horizon Europe Programme under the Specific Grant Agreement No. 101147319 (EBRAINS 2.0 Project). It has also received funding from the European Union’s Horizon Europe Programme under the Specific Grant Agreement No. 101137289 (Virtual Brain Twin Project).

